# Network and systems based re-engineering of dendritic cells with non-coding RNAs for cancer immunotherapy

**DOI:** 10.1101/2020.09.10.287847

**Authors:** Xin Lai, Florian S. Dreyer, Martina Cantone, Martin Eberhardt, Kerstin F. Gerer, Tanushree Jaitly, Steffen Uebe, Christopher Lischer, Arif Ekici, Jürgen Wittmann, Hans-Martin Jäck, Niels Schaft, Jan Dörrie, Julio Vera

## Abstract

Dendritic cells (DCs) are professional antigen-presenting cells that induce and regulate adaptive immunity by presenting antigens to T cells. Due to their coordinative role in adaptive immune responses, DCs have been used as cell-based therapeutic vaccination against cancer. The capacity of DCs to induce a therapeutic immune response can be enhanced by re-wiring of cellular signalling pathways with microRNAs (miRNAs). Since the activation and maturation of DCs is controlled by an interconnected signalling network, we deploy an approach that combines RNA sequencing data and systems biology methods to delineate miRNA-based strategies that enhance DC-elicited immune responses.

Through RNA sequencing of IKKβ-matured DCs that are currently being tested in a clinical trial on therapeutic anti-cancer vaccination, we identified 44 differentially expressed miRNAs. According to a network analysis, most of these miRNAs regulate targets that are linked to immune pathways, such as cytokine and interleukin signalling. We employed a network topology-oriented scoring model to rank the miRNAs, analysed their impact on immunogenic potency of DCs, and identified dozens of promising miRNA candidates with miR-15a and miR-16 as the top ones. The results of our analysis are incorporated in a database which constitutes a tool to identify DC-relevant miRNA-gene interactions with therapeutic potential (www.synmirapy.net/dc-optimization).

## INTRODUCTION

Dendritic cells (DCs) play an important role in regulating adaptive immunity by presenting antigens to T cells (1). Due to the unique function of DCs in the coordination of adaptive immune responses, they have been tested as cell-based vaccination against tumours (2). To obtain immunogenic potency *ex vivo*, monocyte-derived DCs need to go through a complex maturation process, in which DCs are exposed to a monocyte-conditioned medium or a cocktail of cytokines (3, 4). These treatments result in various phenotypic changes in DCs, such as upregulation of co-stimulatory surface markers (e.g., CD80 and CD40) and secretion of pro-inflammatory cytokines (e.g., IL-12 and TNFα). The matured DCs loaded with cancer antigens are infused into patients and trigger a selective immune response by migrating into the peripheral lymphatic tissue, where they encounter and activate tumour-specific T cells (5).

The capacity of DCs to induce an immune response can be improved by molecular engineering. Pfeiffer and co-workers enhanced DCs through electroporation with mRNA encoding a constitutively active variant of IKKβ (caIKK), a kinase upstream of NF-κB that is a key regulator of the immune response (6, 7). Specifically, the kinase phosphorylates IκB resulting in desequestration of the transcription factor (TF) NF-κB and its translocation into the nucleus, where it regulates expression of immune-related genes such as cytokines (8, 9). This engineered IKKβ promotes constant activation of NF-κB signalling, and the cells expressing it (hereafter labelled caIKK-DCs) can induce repeated expansion of Melan-A-specific cytotoxic T cells with a memory phenotype (7). Such DCs are currently under evaluation as vaccine in a phase I clinical trial for the treatment of uveal melanoma patients (NCT04335890).

Our hypothesis is that DCs can be further improved using non-coding RNAs, in particular microRNAs (miRNAs), interacting with key regulators of DC activation and maturation. miRNAs are a class of small endogenous non-coding RNAs with a length of 22-25 nucleotides. Through the inhibition and modulation of the transcription and translation of specific protein-coding genes, miRNAs can alter the basal state of cells and the outcome of stimulatory events (10, 11). Increasing evidence shows that miRNAs play a crucial role in the development and function of DCs (12, 13). They serve as important regulators of complex networks by targeting key signalling genes to regulate proliferation and cell death as well as homeostasis (14). It has also been found that miRNAs are pivotal in both adaptive and innate immunity, e.g., by controlling the differentiation of immune cell subsets and their immunological functions (15). In particular, miRNAs can modulate the immune response by inducing apoptosis, affecting homeostasis, and changing the cytokine profile of DCs (16). Further, one can use miRNAs, alone or in combination, in therapeutic setups to inhibit expression of selected genes in cancer and other targeted cells (17–19).

To facilitate the re-wiring of DCs, it is crucial to understand the intracellular regulatory processes involved in DC maturation and activation. However, the regulatory networks eliciting the activation and maturation of DCs involve multiple interconnected signalling and transcriptional circuits, and their understanding and proper manipulation requires the combined use of high-throughput data and systems biology methods (20, 21). We here present a systems biology approach to understanding the role that miRNAs play in regulating the function of DCs in immunotherapy (Figure 1), and exploit this knowledge to enhance their potential to stimulate an immune response using miRNAs.

**Figure 1.**
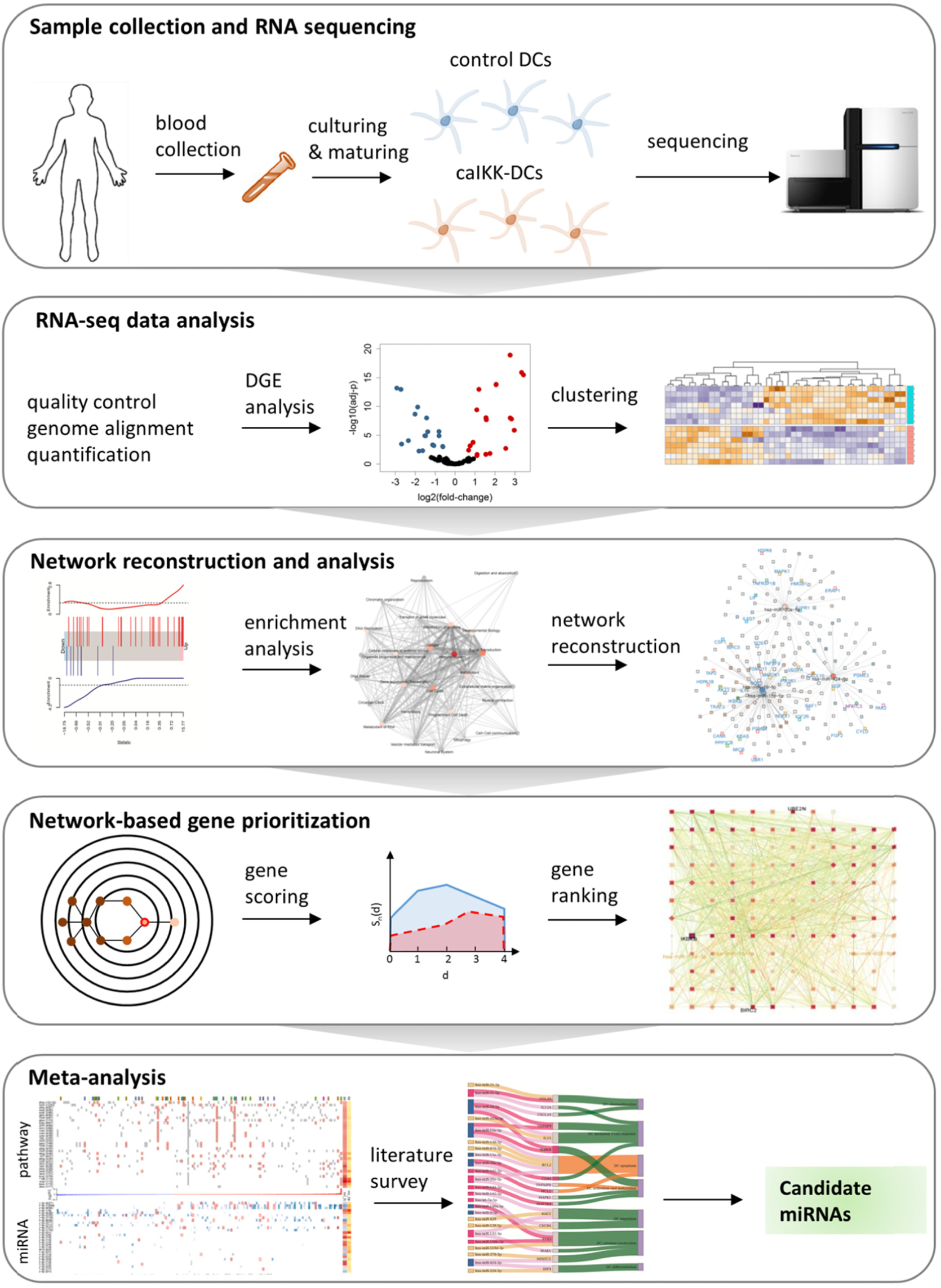
A systems biology approach to study miRNA regulation in DCs. The study starts with preparing of donor DCs followed by cocktail maturation and subsequent electroporation of mRNA encoding Melan-A (control DCs) or a constitutively active variant of IKKβ (caIKK-DCs). The obtained RNA-seq data are processed and analysed for annotating and quantifying protein-coding genes and miRNAs. The identified differentially expressed genes in DCs are used for pathway enrichment analyses and reconstruction of gene regulatory networks. A network topology-oriented scoring model is employed to prioritize miRNAs in different pathway categories of DCs. Finally, a literature review of the top ranking miRNAs in immune signalling pathways elaborates their potential function for improving immunogenic potency of caIKK-DCs.

In this study, we chose the caIKK-DCs as an ideal model system to identify miRNAs that are involved in DC-activation via NF-κB signalling and can boost pro-inflammatory signals. We think the identified miRNAs can enable the DCs to repetitively stimulate T cell expansion. To this end, we performed RNA sequencing (RNA-seq) to obtain the transcriptomic profile (i.e., protein-coding genes and miRNAs) of caIKK-DCs in relation to standard DCs. Next, we analysed miRNA-gene interactions at the pathway level and reconstructed regulatory networks underlying immunological functions of DCs. We then performed network-based prioritization of miRNAs by integrating their expression profiles and their strength of association with other protein-coding genes.

Our analysis identified dozens of miRNA candidates with miR-15a-5p and miR-16-5p as top ones in the regulation of caIKK-DCs. We showed that both miRNAs may exert a strong regulatory effect on genes involved in NF-κB signalling and also target chemokines and cytokines regulating T-cell response. Moreover, we delineated molecular mechanisms through which the miRNAs alter the immunogenic potency of caIKK-DCs. The results of our analysis are available in a web database that facilitates their exploration and visualization (www.synmirapy.net/dc-optimization), thereby providing researchers with a tool to select functional miRNA-gene interactions with therapeutic potential in DCs for experimental investigation.

## MATERIALS AND METHODS

### Generation of monocyte-derived dendritic cells from blood samples

Monocyte-derived DCs were generated as previously described (22). In brief, blood samples from seven healthy donors were collected after approval was granted by the responsible institutional review board (Ethikkommission der Friedrich-Alexander-Universität Erlangen-Nürnberg, Ref. no. 4158) and written informed consent was obtained. Peripheral blood mononuclear cells were isolated from whole blood using density gradient centrifugation. Monocytes were extracted from the non-adherent fraction by plastic adherence, were cultured in DC medium (RPMI (Lonza, Verviers, Belgium) containing 1% non-autologous human plasma (Sigma-Aldrich, St Louis, USA), 2 mM L-glutamine (Lonza), and 20 mg/L gentamicin (Lonza), and were differentiated into DCs by application of 800 U/mL GM-CSF (Miltenyi Biotec, Bergisch Gladbach, Germany) and 250 IU/mL IL-4 (Miltenyi Biotec) on days 1, 3, and 5. After six days in culture, DCs were matured for 24 hours using a cytokine cocktail consisting of 200 IU/mL IL-1β (CellGenix, Freiburg, Germany), 1 000 U/mL IL-6 (CellGenix), 10 ng/mL TNF (Beromun, Boehringer Ingelheim Pharma, Germany), and 1 μg/mL PGE2 (Pfizer, Zurich, Switzerland).

### RNA *in vitro* transcription and DC electroporation

Generation of *in vitro* transcribed RNA with mMESSAGE mMACHINE™ T7 ULTRA transcription-kits (Thermofisher scientific, Waltham, USA) and the electroporation of cocktail-matured DCs (cmDCs) was carried out as previously described (23). For transcriptome analyses, cmDCs were electroporated using 30 µg RNA encoding constitutively active IKKβ (caIKK) since its introduction into mature DCs has been shown to improve activation of T cells (7) and natural killer cells (24). DCs electroporated with RNA encoding melanoma antigen recognized by T cells 1 (Melan-A) were used as control, and such DCs were shown to have no influence on the DCs’ transcriptome profile (25). After electroporation, DCs were cultured in DC medium containing GM-CSF and IL-4 at the concentrations indicated above.

### RNA sequencing processing and differential gene expression analyses

Total RNA including small RNAs was extracted four hours after electroporation using the RNeasy Plus Mini Kit (QIAGEN GmbH, Hilden, Germany) and the generated samples were sequenced using an Illumina HiSeq-2500. Demultiplexed reads were filtered for ribosomal RNAs, transfer RNAs, mitochondrial rRNAs, and mitochondrial tRNAs. The reads were aligned to the human reference genome (hg19) using *STAR* (v2.5.2b) and assigned to genes using *Subread* (v1.5.2). Only uniquely mapping reads that could unambiguously be assigned to a single gene were considered for analysis (Supplementary Table S1).

For miRNA expression quantification, we performed a quality check of the RNA-seq reads with *FastQC* (26), mapped the short sequences to the human reference genome (hg19) using *BWA* (27), and calculated raw read counts of mature miRNAs that are known and annotated in miRBase v21 (28) (Supplementary Table S2; See Supplementary Materials for details).

Before differential expression analysis, we aggregated read counts of Ensembl identifiers that represent the same gene and discarded genes with less than 5 read counts in any sample to increase power for detecting differentially expressed genes (29, 30). Next, we used *DESeq2* (31) in R version 3.6.3 (32) to assess differential expression for protein-coding genes and miRNAs. Then, we performed independent filtering on the results to remove genes that have no or little chance of showing significant evidence (Supplementary Table S3 and S4). Specifically, the independent filtering uses the mean of normalized counts as a threshold to optimize the number of adjusted p-values ≤ 0.05 (31). If the normalized expression of a gene was lower than the threshold, it was discarded. The Benjamini-Hochberg method was then used on the set of remaining genes to correct for multiple comparisons (33). Genes with adjusted p-values ≤ 0.05 were regarded as significantly differentially expressed.

### Gene set enrichment analysis

We extracted all curated pathways from the Reactome pathway knowledgebase (release 68) (34) together with their hierarchical and biological classification according to the database developers. We retraced Reactome’s pathway hierarchy by assigning every pathway from *Homo sapiens* to its corresponding root categories, such as signalling transduction and immune system (see Supplementary Materials for details). As a result, we obtained a table of Reactome pathways matched to the 26 root categories (Supplementary Figure S3 and Supplementary Table S5).

We applied a competitive gene set test to perform gene set enrichment analyses for Reactome pathways. The algorithm CAMERA (35) tests whether the genes in the set are lowly or highly ranked in terms of differential expression relative to genes not in the set, with a positive gene set score indicating a shared tendency for upregulation of the corresponding genes, and *vice versa* (see Supplementary Materials for details). All genes identified as differentially expressed from our RNA-seq data were used as the background gene list for the enrichment analysis. All obtained p-values were corrected using the Benjamini-Hochberg method. Pathways with false discovery rate (FDR) ≤ 0.05 were regarded as significantly up- (positive score) or down-regulated (negative score) in our comparison of caIKK-DCs with controls (Supplementary Table S5). The gene set enrichment analysis was performed using the CAMERA implementation in the package *limma* (36) in R.

### Regulatory network reconstruction

We downloaded functional interactions from the Reactome database (release 68). The collection includes protein-protein interactions, transcriptional regulation, gene co-expression, protein domain interaction, Gene Ontology (GO) annotations and text-mined protein interactions, which cover almost half of the human proteome (37). There are different types of directional molecular interactions including: activation, inhibition or repression, and co-expression or complex formation. The biochemical reactions covered are phosphorylation and ubiquitination. We processed the list to transform bidirectional interactions into their two unidirectional constituents. The result list contained 435,043 unidirectional interactions among 13,852 protein-coding genes.

To derive miRNA-gene interactions, we first obtained conserved and non-conserved miRNA binding sites as predicted by Targetscan version 7.2 (38). Then, we filtered the predicted interactions with experimental evidence from miRTarBase version 8.0 (39) and starBase version 2.0 (40). By doing so, we obtained a list of miRNA-gene interactions (Supplementary Table S6) that contain putative miRNA binding sites with experimental support, such as high-throughput experiments (e.g., RNA-seq, microarray, and Ago CLIP-sequencing) and/or low-throughput experiments (e.g., q-PCR, reporter assay, and Western blot).

The obtained lists of protein-coding genes’ functional interactions and miRNA-gene interactions were used to reconstruct gene regulatory networks from Reactome pathways. Specifically, for a pathway of interest, we built a network around its participating genes and calculated the pairwise Pearson correlation coefficients between interaction partners from their normalized count values in caIKK-DCs. The normalized counts were obtained using the regularized logarithm method (31). In the reconstructed networks, we used the Pearson correlation coefficients to filter out interactions that disagree with their regulation type. We assumed that positive interactions (i.e., activation) require positive Pearson correlations and negative interactions (i.e., inhibition) negative Pearson correlations between interacting molecules. Interactions annotated as gene co-expression or formations of protein complexes were kept, assuming that the involved genes can affect each other’s expression or activity in both directions.

Furthermore, we added annotation to the reconstructed networks’ components in the form of differential expression profiles (i.e., fold-change and FDR), types of genes (e.g., protein-coding gene or miRNA), gene interaction types (e.g., functional interaction or post-transcriptional regulation), gene interaction strengths as denoted by the Pearson correlation coefficients introduced above, and immune categories of genes. Immune categories of protein-coding genes were annotated using curated data from the Immport database (41). Data from the TcoF database were used to identify TFs in our networks (42).

### Gene prioritization in regulatory networks

We prioritized genes in a network using *SANTA* in R (43). The algorithm determines a score of relative importance for each node in a network through a clustering model that accounts for network topology (distances between nodes) and node weights (in our case, a measure of differential expression called perturbation). Briefly, a gene is assigned a high score when itself and its close neighbours in the network have a higher-than-average node weight. The closeness, or distance, between genes is calculated by finding the shortest path through the network.

The node weight is given by the gene’s perturbation (i.e., -log10(adj-p) · |log2(fold-change)|). Both adjusted p-value and log2 fold-change of the gene were taken from the differential gene expression analysis. The distances between neighbouring nodes were calculated as 1 -|*p*|, where *p* represents the Pearson correlation coefficient between the two interacting genes. Higher correlation coefficients (i.e., higher interaction strengths) correspond to lower edge length and thus shorter distance between the nodes. The calculated score was used to prioritize genes (see Supplementary Materials for details). As our networks contain both miRNAs and protein-coding genes that have different types of interactions, miRNAs and protein-coding genes were ranked separately.

### Mapping of microRNA-gene interactions into the curated DC network

To generate the curated DC network, we made use of a previously published network of macrophage pathways (51), since macrophages and DCs are generally considered to be quite similar. We manually added pathways for antigen processing and presentation that were not present in the macrophage map through a comprehensive database search in Reactome. The enriched DC network is a restructuring (see (51) for details on the algorithm) of the curated version that also incorporates miRNA-gene interactions identified () in this study. The reconstructed network is accessible at https://vcells.net/dendritic-cell.

### MicroRNA cooperativity analysis

We used the TriplexRNA database (44) to identify miRNAs that can cooperate with significantly differentially expressed miRNAs in caIKK-DCs to repress protein-coding genes of interest. The obtained RNA triplexes were further filtered using pre-computed equilibrium concentrations and minimum free energies. We kept the RNA triplexes with equilibrium concentrations ≥ 50 nM and minimum free energies ≤ -25 kcal/mol and regarded the participating miRNAs as cooperative partners to repress protein-coding genes.

### Data visualization

The gene regulatory networks for significantly enriched pathways were drawn using *ggraph* (45) and *igraph* (46) in R. Heat maps were plotted using *Complexheatmap* (47) in R. Scatter and bar plots were drawn using *ggplot2* (48) in R. Sankey diagram was drawn using *networkD3* (49) in R. The clustered Reactome pathways were visualized using Cytoscape version 3.72 (50).

## RESULTS

### Transcriptome analysis reveals miRNA expression changes in caIKK-DCs

To characterize the gene expression profiles of DCs treated by caIKK, we collected blood samples from seven healthy donors and generated monocyte-derived DCs (see Materials and Methods). Each of the matured DC cultures was split for the subsequent experiment: One half was electroporated with mRNA encoding constitutively active IKKβ (caIKK, encoded by an engineered *IKBKB*) and the other half with mRNA encoding the melanoma antigen Melan-A (encoded by *MLANA*). The latter was used as control, as maturated DCs transfected with Melan-A mRNA did not show any significant changes in their transcriptomic profiles compared to untreated DCs (25). Our data showed that caIKK electroporation induced maturation and activation in DCs, including secretion of pro-inflammatory cytokines such as IL-6, IL-8, IL-12, and TNFα (Supplementary Figure S1A), upregulation of co-stimulatory surface markers such as CD25, CD40, CD70, CD80, CD86, and OX40L (Supplementary Figure S1B), and increased maturation as indicated by the expression of CD25 and CD70 in the seven donors (Supplementary Figure S1C).

Four hours after electroporation, RNA was isolated and assessed via bulk RNA sequencing (RNA-seq). We chose this early time point because we were interested in mRNA levels, which are expected to quickly respond to the activation of NF-κB as a result of continuous IKKβ activation. From the RNA-seq data, we identified 63 protein-coding genes and 44 miRNAs that were significantly differentially expressed (DE) between caIKK-DCs and controls (Supplementary Table S3 and S4; see Materials and Methods). Among the protein-coding genes, *MLANA* (encoding Melan-A) and *IKBKB* (encoding IKKβ) were the most down- and upregulated in caIKK-DCs, respectively (Supplementary Figure S2A). This is in line with the fact that the mRNA content of the two genes was artificially altered in the respective populations and can be considered a quality control for the experimental results. For the miRNAs, miR-146a/b and miR-155 were upregulated in caIKK-DCs, in consistence with them being transcriptional targets of NF-κB (52) and being upregulated in mature DCs (53). By performing principal component analysis, we assessed the clustering tendency in the RNA-seq data. Controls and caIKK-DCs showed better separation when restricting the input to the measured miRNAs rather than the whole transcriptome (Supplementary Figure S2B). In addition, the DE miRNAs unequivocally separated the caIKK-DCs from the controls in hierarchical clustering (Supplementary Figure S2C). These results suggested that caIKK-DCs harbour a distinct miRNA expression profile.

### The gene signature induced in caIKK-DCs is associated with NF-κB activation

To understand the molecular function of the identified DE genes in the caIKK-DCs, we performed gene set enrichment analysis using the Reactome pathway database. The database contains more than 2,000 cellular pathways curated from 30,721 peer-reviewed publications and classified into 26 root categories (34), thereby enabling a systematic and comprehensive analysis of DE genes. The 26 categories consist of a set of pathways that are annotated to be hierarchically and functionally linked. We calculated enrichment scores for each pathway which reflect the degree to which its corresponding gene set tends to be up- or downregulated in caIKK-DCs (Figure 2A; see Materials and Methods).

**Figure 2.**
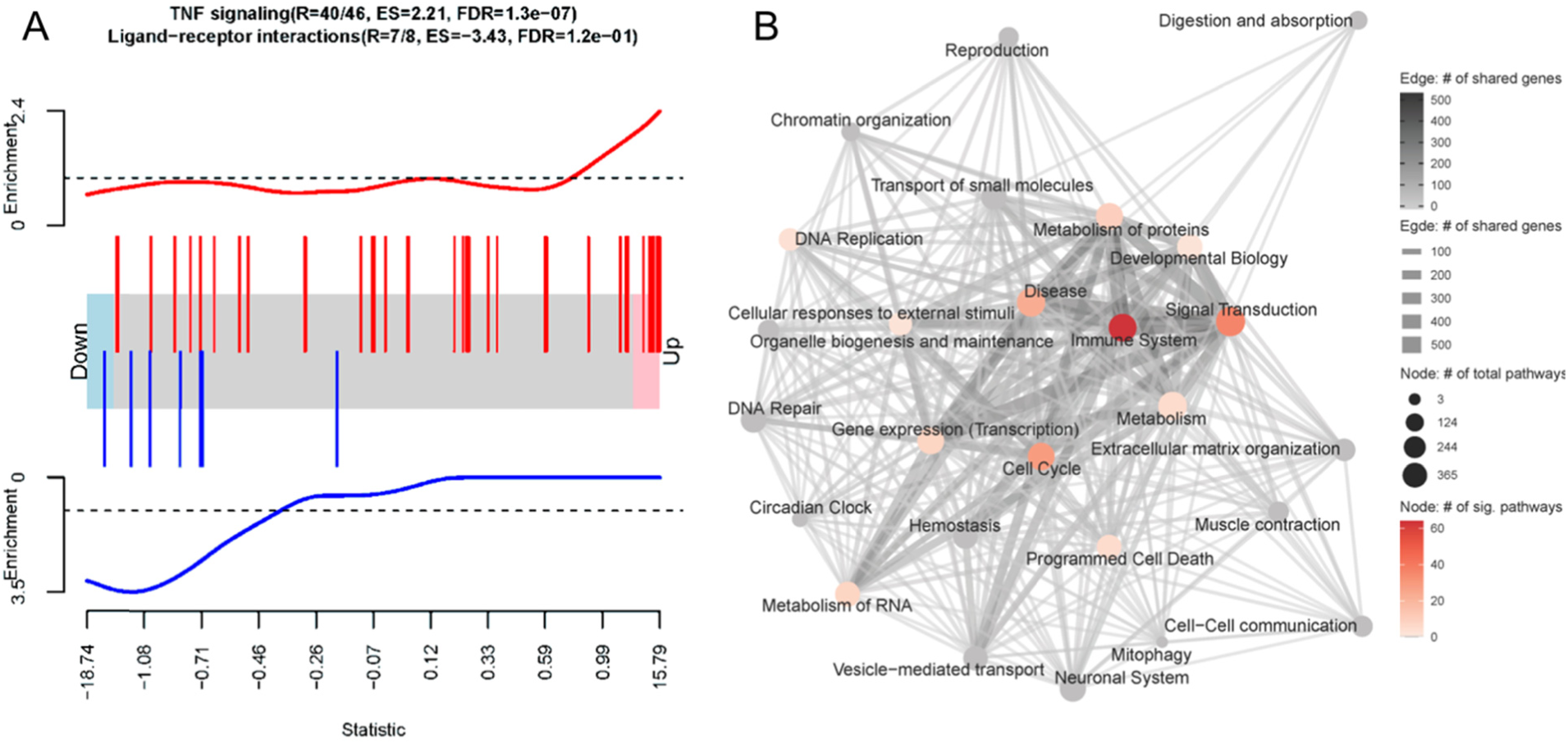
Gene set enrichment analysis on the differential expression analysis between caIKK-enhanced and control DCs. (**A**) Enrichment plots of two exemplary gene sets. The genes’ weighted log2 fold-change (i.e., log2 fold-change divided by the standard error of the log2 fold-change) obtained from the differentially gene expression analysis are sorted from the smallest to the largest (the barcode plot). Genes from *TNF signalling* (red bars) accumulate at the upregulated end while genes from *ligand-receptor interaction* (blue bars) in Hedgehog signalling accumulate at the downregulated end. The curves show the local enrichment score of the vertical bars in the barcode plot. For the red curve, parts above the dashed line signify enrichment while parts below the line signify depletion. For the blue one, parts below the dashed line signify enrichment while parts above the line signify depletion. (**B**) Network visualization of the gene set enrichment results for Reactome’s 26 root categories (nodes annotated with texts). Categories are connected when they have shared genes (edges). The size of a node denotes the number of pathways that belongs to the respective category. The node colour represents the number of significantly enriched pathways in a category. A grey node means that no significant pathways were identified in the category. The width and colour of an edge represent the number of shared genes between the two connected categories. A detailed map containing all pathways and their corresponding enrichment scores can be found in Supplementary Figure S3.

We found that the DE genes in caIKK-DCs are significantly enriched in 195 Reactome pathways, most of which belonged to the Reactome categories *signal transduction* and *immune system* (Figure 2B; Supplementary Figure S3 and Table S5). In the category of *immune system*, 65 out of 182 pathways were identified as significantly enriched, including cytokine signalling pathways and pathways associated with innate and adaptive immune response. This suggested that the continuous activation of IKKβ in DCs has a generalized effect on DC-mediated immune responses. In addition, we identified 12 enriched pathways that are directly associated with NF-κB activation and signalling (see Supplementary Table S5), such as NF-κB activation by IκB kinase complexes. This was consistent with the current model of the canonical NF-κB activation pathway, in which IKKs phosphorylate IκB resulting in desequestration of NF-κB and its translocation into the nucleus, where it regulates expression of immune-related cytokine genes and others(8, 9). All these NF-κB pathways had positive enrichment scores, indicating that the genes involved are more likely to be up-regulated in the caIKK-DCs. The results were in line with our expectation that the caIKK-DCs can trigger a stronger immune response as a result of NF-κB desequestration by constitutive activation of IKKβ.

### Significantly differentially expressed miRNAs in caIKK-DCs regulate an abundance of enriched immune pathways by targeting hundreds of their protein-coding genes

To identify potential miRNA-gene interactions regulating the immunogenic potency of DCs, we first obtained putative miRNA-gene interactions for the significantly DE miRNAs (see Materials and Methods). We kept the putative interactions that are validated by experiments. For each identified miRNA-gene interaction, we then computed the Pearson correlation coefficient between miRNA and target gene expression. The interactions with negative correlation were regarded as reliable and functional, as miRNAs canonically repress translation initiation or stimulate mRNA degradation (54) and miRNA-mediated gene activation usually results from indirect regulation mechanisms (55). The data showed that 36 out of the 44 miRNAs are involved in the regulation of protein-coding genes belonging to 195 enriched pathways of the 26 Reactome root categories (Figure 3A).

**Figure 3.**
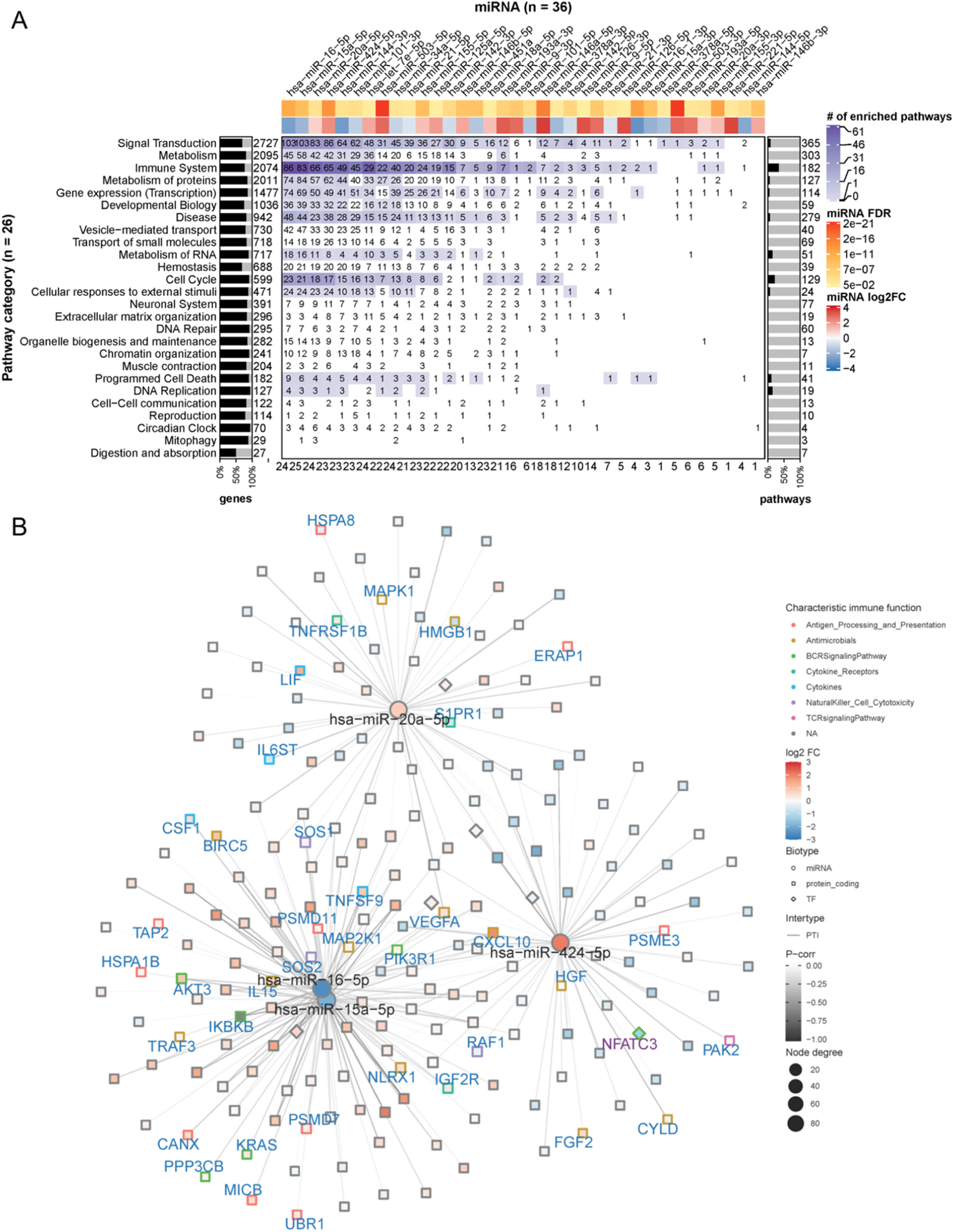
miRNA targeting profiles in Reactome pathway categories. (**A**) Overview of the 36 DE miRNAs (in columns) targeting the 26 Reactome pathway categories (in rows). On the heat map grid, the number of protein-coding genes targeted by the respective miRNA is given, and the colour represents the category’s number of significantly enriched pathways that are regulated by the miRNA. For example, if an entry shows *n* with a colour corresponding to *m* on the figure legend, it means a miRNA regulates *n* targets in *m* significantly enriched pathways of a category. The bar plot on the left indicates the per-category percentage of the protein-coding genes (black bars) that were found in our RNA-seq data, with the total number of molecules in the category given next to it. The bar plot on the right indicates the percentage of enriched pathways (black bars) per category, with the total number of pathways in the category given next to it. The figures at the bottom tabulate how many categories a miRNA regulates. The top annotation shows statistics from the differential expression analysis for the miRNAs (i.e., fold-change in log2 scale and FDR). (**B**) The network shows miRNA-gene interactions in the category *signal transduction*. The four miRNAs (miR-16, miR-15a, miR-20a, and miR-424) that have the largest number of targets were selected. The node size is proportional to the node degree. The node colour represents a gene’s fold-change in log2 scale. The node shape denotes the type of a gene, including protein-coding (square) and miRNA (circle), with their names shown in blue and black labels, respectively. TFs are drawn as diamonds with their names shown in purple font. The colour of node borders represents different categories of annotated immune genes, with the gene names given as labels. The edge colour shows Pearson correlation coefficients between the expression of miRNAs and that of their targets.

In individual pathway categories, the number of molecules (i.e., protein-coding genes, DNA/RNA, drugs, and chemical compounds) ranged from 27 to 2727, and genes identified by our RNA-seq data covered between 51% and 95% of the molecules in the respective category. For all categories except *digestion and absorption*, the DE miRNAs were identified to regulate 1 to 103 protein-coding genes (denoted by the numbers on the heat map grid cells in Figure 3A). Some miRNAs were found to regulate the expression of dozens of protein-coding genes in more than ten categories, suggesting that they act as regulatory hubs in caIKK-DCs. For example, miR-15a-5p, miR-16-5p, miR-20a-5p, and miR-424-5p can potentially regulate more than 80 protein-coding genes in the category *signal transduction* (Figure 3B), and they also have more than 60 targets in *immune system*. In contrast, miR-15a-3p and miR-9-3p target only BCL2 and MAPK1 in *immune system*, suggesting a specific role for them in regulating cytokine signalling in caIKK-DCs (Supplementary Figure S4). Furthermore, we found that some miRNAs target a high fraction of enriched pathways belonging to specific Reactome root categories (denoted by the colour of heat map grid cells in Figure 3A). For instance, miR-16-5p regulates 61 out of 64 enriched *immune system* pathways and 34 out of 36 enriched pathways in *signal transduction*. This suggested that it plays a vital role in regulating the immunogenic potency of caIKK-DCs. On the other hand, some categories included abundant enriched pathways that are regulated by multiple DE miRNAs. Interesting cases were pathways associated with *protein metabolism, RNA metabolism, programmed cell death*, and *cell cycle*. This result suggests that the DE miRNAs in caIKK-DCs are involved in regulating synthesis, processing, and modification of mRNAs and proteins and can also participate in other biological processes, such as cell cycle and cell apoptosis. Taken together, the DE miRNAs in caIKK-DCs target and potentially coordinate the activity of immune-relevant pathways in a pleiotropic fashion.

### Network-based prioritization of miRNAs in caIKK-DCs

The ubiquitous, pleiotropic, and concerted gene regulation by miRNAs makes it challenging to quantify the relative impact of each individual miRNA. To prioritize the DE miRNAs according to their potential to act synergistically with NF-κB in DC activation, we applied a network-based method that integrates their expression and interaction profiles.

First, we reconstructed one gene regulatory network for each of the 26 pathway categories. The reconstructed networks were composed of miRNA-gene interactions and functional interactions among protein-coding genes. Interactions were discarded when the sign of their Pearson correlation coefficient of expression disagreed with their regulation type, such as inhibition or activation (see Materials and Methods). Depending on the category, the size of the corresponding networks varied from 1,915 genes and 57,520 interactions (for *signal transduction*) to 30 genes and 153 interactions (for *mitophagy*). To prioritize the network components involved in regulating the immunogenic potency of DCs, we used a clustering model (43) to calculate a node score (Figure 4A; see Materials and Methods).

**Figure 4.**
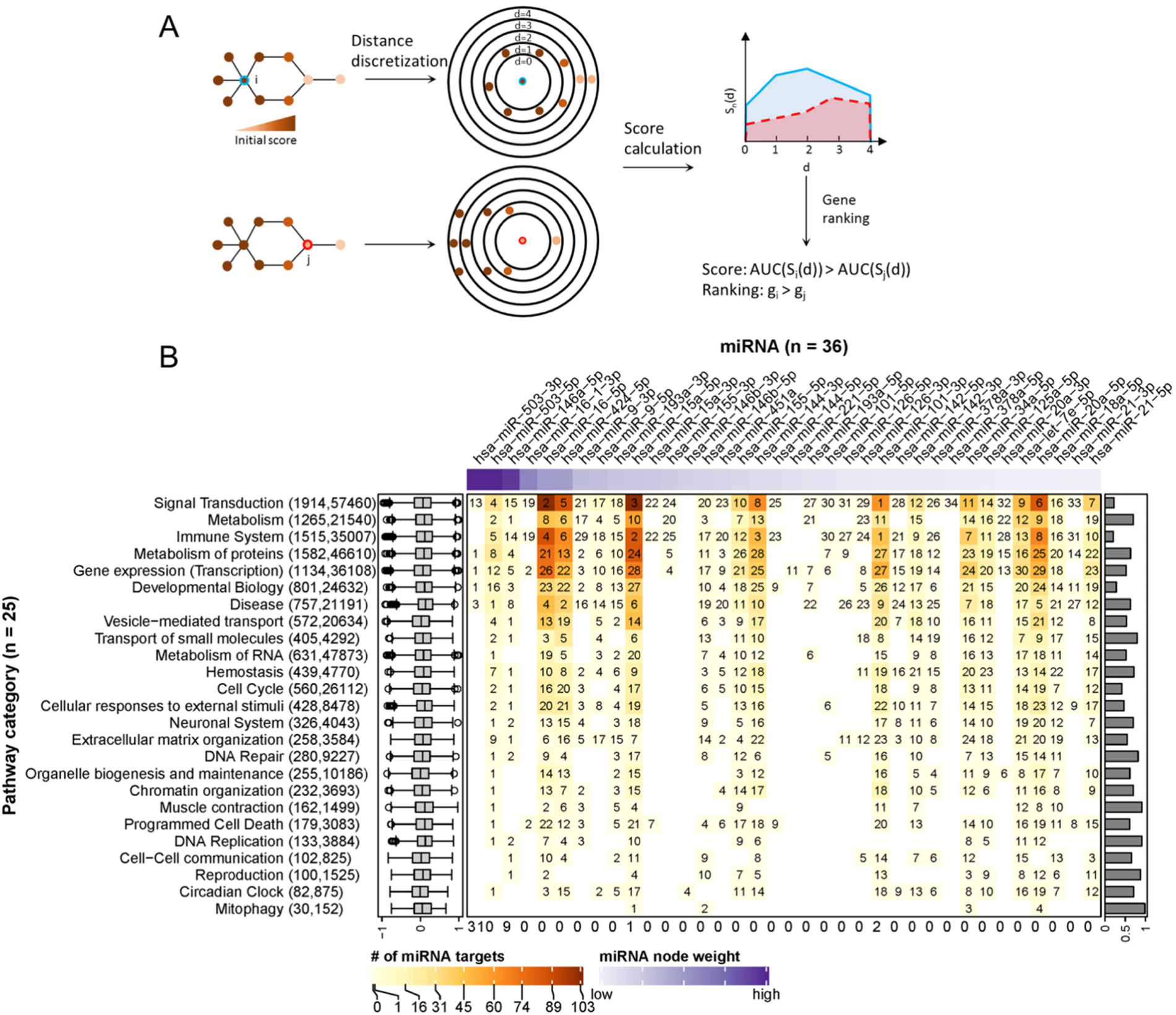
Ranking of miRNA relevance based on expression and regulatory network neighbourhood. (**A**) Network nodes were scored with an algorithm that uses the guilt-by-association principle to rank genes. In other words, a gene inside of or close to a cluster of important genes is potentially more important than a gene that is further away. In our case, the importance of a gene for a phenotype is quantified by their perturbation in expression (denoted by node colours). In a network, the distance of the gene in question (red or blue border) to other genes is calculated via the weighted shortest-path method. The range of observed distances is then subdivided into discrete bins (denoted by circles in the figure) and an estimate of neighbourhood importance calculated for each bin, i.e., for all nodes up to the respective distance. The area under the curve (AUC) in the plot of bins (*d*) vs neighbourhood importance is used as the gene’s score. In the example, gene *i* is much closer to genes with high node weights (i.e., their perturbation in caIKK-DCs is large) than gene *j*. As a result, the blue area is bigger than the red area, and thus gene *i* ranks higher than *j*. (**B**) Heat map of miRNA ranking in pathway categories. The columns of the matrix indicate the 36 DE miRNAs sorted by perturbation (i.e., node weights), and the rows of the matrix indicate 25 categories of Reactome pathways. The category *digestion and absorption* is not shown, as we did not identify functional and reliable interactions among its genes. On the heat map grid, the rank of a miRNA in a category is given as a number, and the colour represents the number of protein-coding genes targeted by it. For example, if an entry shows *k* with a colour corresponding to *n* on the figure legend, it means that the miRNA ranks *k*th (1st is the highest ranking) and regulates *n* targets in the category; A white grid cell means that the miRNA has no targets in the category and thus no ranking. The top annotation shows node weights of the miRNAs. The numbers in parentheses on the left side list how many genes and edges the reconstructed regulatory network of the category possessed. The box plots on the left show the distribution of edge weights (denoted by Pearson correlation coefficients between genes) in the networks. The bar plots on the right show the Pearson correlation coefficients between a miRNA’s perturbation and its score. The numbers at the bottom show the times miRNAs ranked 1st in the pathway categories.

The score ranked *IKBKB*, whose expression was greatly increased by mRNA electroporation, as the top protein-coding gene in 51 out of 59 networks in which it is involved (Supplementary Table S7; Supplementary Figure S5). This result is consistent with our expectation that the intentionally modulated gene in experiments is prioritized, and thus demonstrating the ability of the model to identify crucial regulatory genes in the experiments. In the two prominent categories *signal transduction* and *immune system*, the NF-κB family and genes related with immune signalling or antigen processing and presentation tended to rank higher than other genes (Supplementary Figure S6). This result again justified the ability of the model to prioritize important genes in networks, as members of the NF-κB family are downstream targets of *IKBKB* while signalling and antigen presentation genes are supposed to be crucial regulating immune function of DCs.

Furthermore, we analysed the data to identify crucial miRNAs for each Reactome root category. As shown in Figure 4B, miRNAs with higher node weights (i.e., stronger perturbation) generally ranked higher in a category, as miRNA scores and node weights showed a positive correlation, ranging from 0.19 to 0.98. Specifically, miR-503-5p, miR-503-3p, and miR-146-5p had the highest perturbation in the DE miRNAs, and they ranked top in 22 out of 26 categories. However, the interaction profile also plays a role, as for example in *signal transduction*, the three top-ranking miRNAs miR-101-3p, miR-16-5p, and miR-15a-5p had lower perturbation than miR-146-5p but interacted with more protein-coding genes. In addition, the three above miRNAs and miR-144-3p ranked top in *immune system*, most probably due to the reason that they regulate a large number of protein-coding genes associated with immune signalling pathways. To facilitate the visualization of our results, we integrated the data and the identified miRNA-gene interactions into a comprehensive, manually curated regulatory network including key pathways in DC priming and activation according to the literature (see Materials and Methods; https://vcells.net/dendritic-cell).

Taken together, the reconstructed regulatory networks underlying different cell functions allowed us to identify important miRNA regulators based on their expression and interaction profiles. The miRNAs with the highest scores possibly exert regulatory functions, and manipulation of their expression levels may enhance the immunogenic potency of DCs.

### Potential miRNA-gene interactions to improve caIKK-DCs

To characterize the functional role that miRNAs play in caIKK-DCs, we delineated landscapes of miRNA-gene interactions in the significantly enriched pathways that were found in corresponding categories (see Figure 5 and www.synmirapy.net/DC-optimization). The interaction landscapes are a way of systematically mapping relevant gene interactions, and in our case they served as a tool for identifying functional miRNA-gene interactions in DC priming and activation. As we were particularly interested in identifying miRNAs that can enhance the caIKK-DCs’ immunogenic potency, we focused on analysing miRNA-mediated gene regulation in the category *immune system*. In this category, we identified hundreds of miRNA-gene interactions in significantly enriched pathways, including toll-like receptor, cytokine, and interleukin signalling as well as MHC processing and presentation. All of these pathways had positive enrichment scores, indicating that the involved genes tended to be upregulated in caIKK-DCs according to our analysis. Most protein-coding genes used as indicators of DC activation and maturation (2, 53, 56, 57) were found to be upregulated in the enriched pathways (Supplementary Table S8). The activation of NF-κB signalling led to upregulation of surface proteins that can prime T cells (e.g., *CD40, CD70, CD80*, and *CD86*), chemokines (e.g., *CCL3* and *CXCL10*) that are necessary for T-cell migration, TNF superfamily members that can induce crosstalk between T cells and DCs (e.g., *TNF, TNFRSF4*, and *TNFSF9*), and cytokines that are responsible for stimulating proliferation and activation of T cells (e.g., *CXCL8, IL6, IL12A*, and *IL12B*).

**Figure 5.**
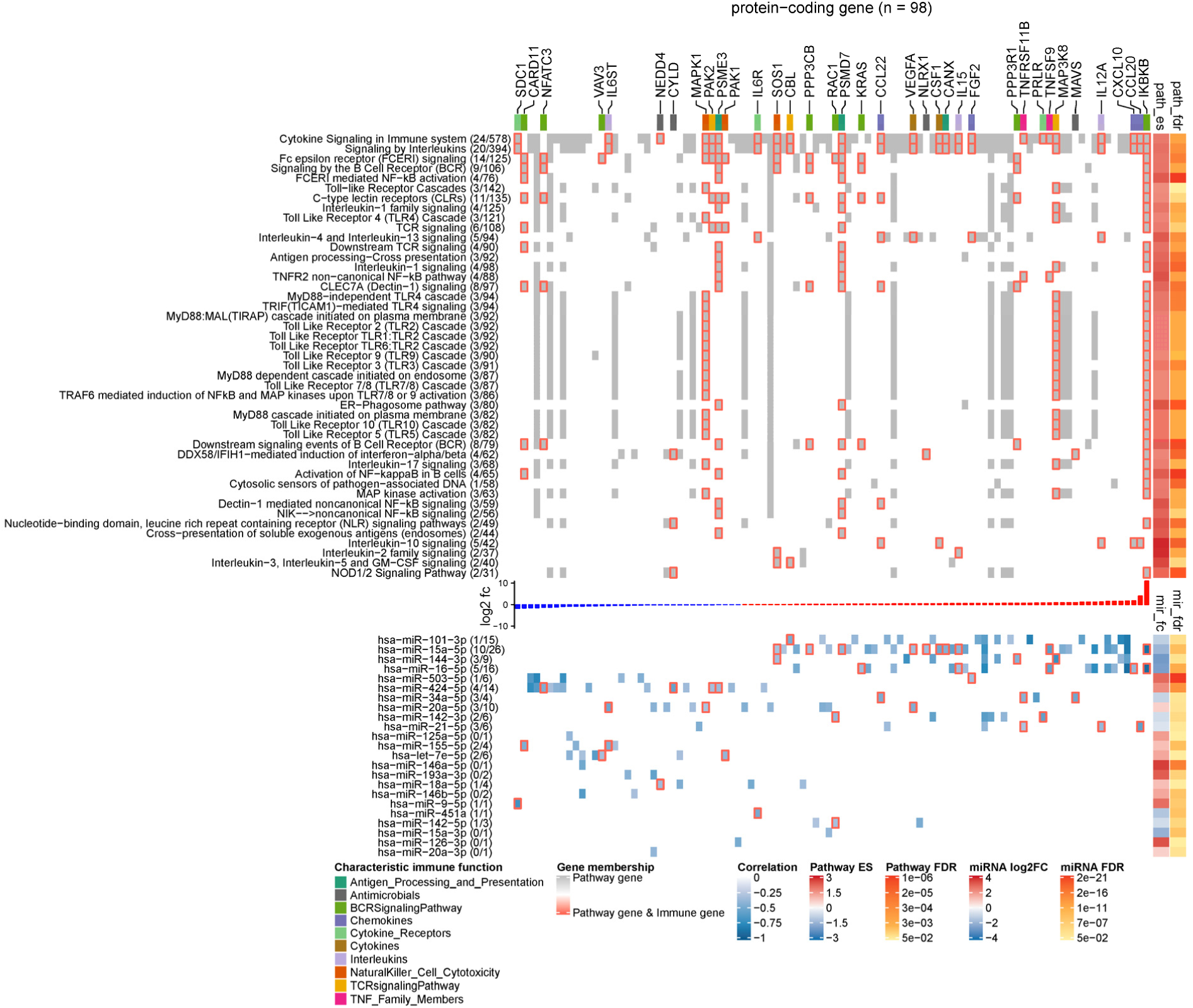
Landscape of miRNA-mediated DC gene regulation in immune signalling pathways. The heat map has two components that share a set of columns corresponding to 98 DE protein-coding genes that are targeted by the DE miRNAs. In the upper component, rows represent pathways from the category *immune system*, and grid cell colours indicate whether a protein-coding gene is involved in the pathway (grey), involved in the pathway and an immune gene (grey grid cells with red borders), or not involved in the pathway (white). The figures next to a pathway name indicate how many DE immune genes (left) and how many protein-coding genes found in our RNA-seq data (right) belong to it. The top annotation highlights genes with different characteristic immune function using a colour code. The annotation on the right side shows the statistics of the gene set enrichment analysis including the enrichment score and the FDR. The bar plot between the heat map components shows the log2 fold-change of the genes in caIKK-DCs (blue: downregulated; red: upregulated). In the lower component, the rows represent the ranking miRNAs in *immune system* (from high to low) and the grid cells show the regulative influence of a protein-coding gene by a miRNA, which is estimated by the Pearson correlation coefficients between their expression profiles. If a gene is a known immune gene, the corresponding grid cell has a red border. The numbers in the parentheses next to the miRNA names show the number of DE immune genes and the number of DE protein-coding genes that are regulated by a miRNA. The right annotation shows the results of the differential expression analysis including the log2 fold-change of miRNA expressions and their FDRs. For lack of space, we show only enriched pathways with more than 30 protein-coding genes picked up in the RNA-seq data, and in each pathway, only a subset of protein-coding genes that are estimated to be strongly influenced by the miRNAs (Pearson correlation *≤* -0.3) are shown. The complete landscape of miRNA-gene interactions in *immune system* is shown in Supplementary Figure S7.

Furthermore, our data showed that the identified DE miRNAs have a regulative influence (represented by Pearson correlation ≤ -0.3) on protein-coding genes associated with NF-κB activation, cytokines, chemokines, and TFs that are associated with immunophenotypes of DCs (Figure 6). Some of the DE miRNAs were found to cooperate with other miRNAs to regulate the expression of a protein-coding gene (see Materials and Methods). This mechanism, known as miRNA cooperativity, is characterized by more efficient inhibitory effects on the target’s expression compared to the regulation by individual miRNAs (17– 19). Moreover, for most of the identified miRNAs, our analysis proposed specific modulation of their expression levels to improve immunogenic potency of DCs. However, in some cases, such as miR-34a-5p and miR-20a-5p, up- or downregulating their expression levels may result in contradictory effects in DC-mediated immune response, thereby requiring further analysis and experimental investigation. In the following paragraphs, we illustrate and discuss specific functions of the miRNAs in caIKK-DCs.

**Figure 6.**
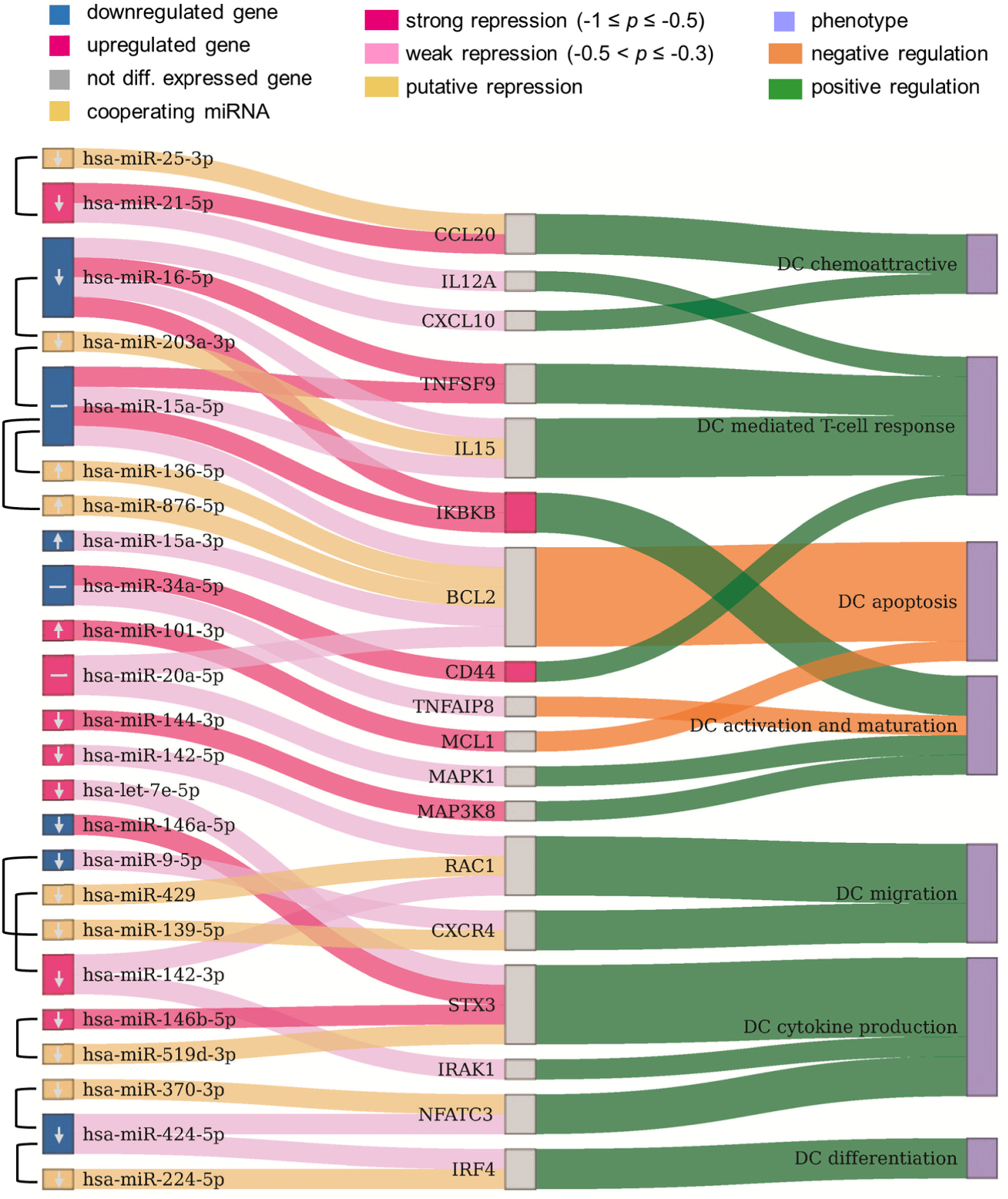
Potential miRNAs for improving immunogenic potency of caIKK-DCs. The Sankey diagram contains three columns that are nodes representing the DE miRNAs from our RNA-seq data and their cooperating miRNAs, protein-coding genes targeted by the miRNAs, and DC phenotypes associated with the protein-coding genes. Two miRNAs that were identified to cooperatively repress a protein-coding gene are connected by a line. Colours of miRNAs and protein-coding genes indicate whether or not they are significantly DE in caIKK-DCs. Arrows in miRNA nodes indicate manipulation of their expression levels to improve immunogenic potency of caIKK DCs (↑: upregulation; ↓: downregulation; −: both direction due to the pleiotropic nature of miRNA in regulating gene expression). Connections between miRNAs and protein-coding genes show regulative influence of protein-coding genes by miRNAs (strong: Pearson correlation ≤ -0.5; weak: -0.3 ≤ Pearson correlation < -0.5). The connections between protein-coding genes and phenotypes denote how a gene regulates a phenotype, and such information was obtained from literature. For instance, miR-424-3p and miR-224-5p target the IRF4 that is known to positively regulate differentiation of DCs. The two miRNAs cooperatively repress the protein-coding gene, and the downregulated miR-424-3p exerts a weak inhibitory effect on the expression of IRF4. A detailed discussion of the results can be found in the main text. The corresponding miRNA-gene interactions in *immune system* and annotated gene-phenotype associations can be found in Supplementary Table S9 and Table S10, respectively.

Chemokines direct cell migration via induction of chemotaxis. For example, CCL5 and CXCL10 improve CD8+ T-cell infiltration (58) and CCL20 plays a role in recruiting regulatory T cells and T helper (Th) 17 cells (59, 60). CXCL10 is repressed by miR-16-5p and CCL20 is targeted by miR-21a-5p with its cooperating miR-25-3p. This suggested that the miRNAs exert an inhibitory function in the recruitment of T cells. miR-21a-5p also targets IL12A, a subunit of the inflammatory cytokine IL-12 that is necessary for CD8+ T-cell clonal expansion, function and memory (56, 61).

Control of DC survival is necessary for maintaining their homeostasis (62, 63). We showed that miR-15a-5p with its cooperating miRNAs (i.e., miR-156-5p and miR-876-5p) and miR-20a-5p target BCL2, and miR-101-3p targets MCL1. The repression of the BCL2 family of anti-apoptosis genes by these miRNAs suggested their ability to undermine the survival mechanism of DCs.

miR-424-5p and miR-224-5p can co-repress IRF4, which is a member of the interferon-regulatory family and can regulate differentiation of specific DCs that can induce Th 2 cell responses (64). miR-20a-5p and miR-144-3p regulate the MAPK signalling pathway by targeting MAPK1 and MAP3K8 respectively, and these MAP kinases have been found to activate the IKK complex that triggers NF-κB activation (65, 66) and also to regulate release of TNFα by DCs (67). miR-34a-5p has a strong regulative influence on CD44 whose presence is important for the immune synapse between DCs and T cells that subsequently regulates T-cell activation (68) and apoptosis (69). miR-34a-5p also targets TNFAIP8 whose knockdown in DCs has been found to promote DC maturation and activation followed by increased proliferation and differentiation of T cells (70). miR-9-5p can cooperate with miR-139-5p to repress CXCR4 that is required for DC migration into the skin’s draining lymph nodes (71). miR-142-3p with its cooperating miR-429 and miR-142-5p target the small GTP-binding protein RAC1 that controls the formation of dendrites in mature DCs and their migration toward T cells (72).

Some identified DE miRNAs target protein-coding genes involved in regulating the DC-mediated secretion of cytokines that are important for the T-cell response. The repression of NFATC3 by miR-424-5p and its cooperating miR-370-3p suggested a regulating influence on the production of IL-2 that is involved in T-cell priming (73, 74). STX3 has been shown to play a role in trafficking of IL-6 or MIP-1α in DCs and thus regulating their secretion (75) and is targeted by let-7e-5p, miR-146a-5p, and miR-146b-5p with its cooperative partner miR-519d-3p. The deficiency of IRAK1 in plasmacytoid DCs abrogates IFNα production, leading to a remodulation of T cell function (76–78), and IRAK1 is a target of miR-142-3p.

Finally, miR-16-5p and miR-15a-5p can cooperate with miR-203a-3p to repress IL-15, an interleukin which can induce T-cell proliferation, enhance cytolytic effector cells including natural killer and cytotoxic T cells, and reinforce B-cell stimulation (79). A recent *in vivo* study has shown that an IL15-enhanced DC vaccine is a potent delayer of tumour growth, improves mouse survival, and induces a stronger Th1-skewed T-cell response (80). The two miRNAs also target the receptor TNFSF9 (also known as CD137), whose stimulation in DCs by its ligand CD137L can lead to secretion of IL-6 and IL-12 and induce T-cell proliferation (56, 81, 82). In addition, miR-16-5p and miR-15a-5p were identified to strongly repress IKBKB itself. Since both miRNAs were found to be downregulated in caIKK-DCs, this implied a positive feedback loop in NF-κB signalling as the miR-15/16 cluster is a transcriptional target of NFKB1 (83). The results suggested both miRNAs as promising candidates for improving the immunogenic potency of caIKK-DCs, as they not only have the ability to strengthen NF-κB activation but also to improve DC-induced immune responses through regulating cytokines and chemokines.

## DISCUSSION AND CONCLUSIONS

We applied a systems biology approach to investigate the regulatory functions of miRNAs in caIKK-DCs. Due to the promiscuous binding of miRNAs, it is challenging to identify relevant miRNA-gene interactions for experimental validation and cell re-engineering (84, 85). Our approach, which integrates transcriptomic profiling, networks of curated signalling pathways, and a prioritisation score, allows the systematic identification of condition-specific miRNA-gene interactions.

Through RNA sequencing of monocyte-derived DCs matured with a cytokine cocktail and electroporated with caIKK mRNA, we identified DE protein-coding genes and miRNAs in the caIKK-DCs. The identified DE miRNAs correctly separated the caIKK-DCs from the control, suggesting a well-defined transcriptional response to caIKK that is consistent with our understanding that miRNAs act as post-transcriptional regulators of expression in DC differentiation and function (86). Among the identified miRNAs there were several, such as miR-15a-5p, miR-16-5p, miR-20a-5p, and miR-424-5p, which target a considerable number of genes. Such hubs have been shown to be important regulators, as they represent sites of signalling convergence in gene regulatory networks and coordinate cell development and function (87– 89). In contrast, other DE miRNAs, such as miR-15a-3p and miR-9-3p, exert a narrow function by regulating the expression of specific protein-coding genes in the caIKK-DCs.

Integration of the transcriptomic response into the curated pathways from Reactome provides an understanding of the functional changes at the pathway level. The gene set enrichment analysis highlighted cytokine, interleukin, and toll-like receptor signalling pathways that are involved in regulating various aspects of innate and adaptive immune responses (90). Such results may be compromised when other pathway databases such as KEGG (91) and WikiPathways (92) are employed, as the relevant pathways and molecular interactions in the pathways are different from Reactome (93, 94). To circumvent this issue, one possibility could be to extract the overlapped networks between the different databases of pathways; however, this is not always possible due to the differences in annotation of genes and interactions. An alternative option is to integrate the data and the detected miRNA-gene interactions into comprehensive, manually curated regulatory networks based on the current literature on DC regulation. This way, one can put the newly discovered relevant interactions into the context of the existing knowledge and facilitate the mining and interpretation of the omics data (95, 96). However, when used inappropriately, knowledge-based networks mainly rediscover existing knowledge but may overlook insights gained from the evaluation of an all-encompassing network.

We reconstructed regulatory networks from Reactome pathways and used them to rank genes and miRNAs according to their predicted impact on DC function. Systematic computation of such a ranking supports and facilitates experimental efforts, allowing them to focus on the most promising candidates. Gene prioritization algorithms have been widely used in recent times to rank genes in networks (97, 98). For instance, the PageRank algorithm designed to analyse the relative importance of websites was adapted to identify crucial genes in biological networks (99), and diffusion-based methods were used on dense networks to prioritize genes (100). We used a gene prioritization algorithm that utilizes the guilt-by-association principle to rank genes based on their own perturbation, i.e., differential expression profile, and their weighted distances to other perturbed genes in a network. The algorithm prioritized dozens of miRNAs, of which miR-16-5p and miR-15a-5p are the top candidates to regulate the immunogenic potency of DCs.

Finally, an in-depth analysis of the identified miRNA-gene interactions in immune signalling pathways showed diverse roles of the DE miRNAs in regulating DC-mediated immune response. For instance, miR-16-5p and miR-15a-5p may have strong regulatory influence on IKBKB that activates NF-κB and on TNFSF9 that controls cytokine secretion of DCs; both miRNAs could have weak inhibitory effects on BCL2 that maintains DC homeostasis and on cytokines (such as CXCL10 and IL15) that regulates T-cell response but may cooperate with other miRNAs to more efficiently repress the protein-coding genes. While these predictions were made based on validated miRNA binding sites and negative correlation between the expression levels of miRNAs and their targets, they cannot quantify the strength of individual repression effects (101). The results suggested both miRNAs as potential candidates for improving immunogenic potency of caIKK-DCs through strengthening NF-κB stimulation and also synergistically regulating other genes related with immunogenic potency.

For most identified crucial miRNAs, our analysis suggested up- or downregulation of their expression levels to improve immunogenic potency of DCs, but in some cases the pleiotropic nature of miRNAs in regulating gene expression makes it difficult to decide how to experimentally modulate their expression. In addition, it is worth noting that the results reflect the early transcriptional response that may differ from that in the long-term. From an experimental perspective, the next step would be to analyse the kinetics of the expressions of miRNA and mRNA after the activation of NF-κB. Further, it remains to be tested how co-electroporating the selected miRNAs, or artificial antagonists thereof, with caIKK or introducing them into the cells after a delay will alter the DCs’ phenotype and immunogenic potency.

Taken together, our approach enables the systematic analysis and identification of functional miRNA-gene interactions that can be experimentally tested for improving DC immunogenic potency. Additionally, since the approach is not specific for DCs, it can be adapted to study miRNAs in other immune cells and relevant immunotherapies.

### DATA AVAILABILITY

The RNA-seq data of DCs is deposited in ArrayExpress.

The database for selecting functional miRNA-gene interactions with therapeutic potential in DCs for experimental investigation is available at: www.synmirapy.net/DC-optimization.

The curated DC network containing identified miRNA-gene interactions and the RNA-seq data are available at https://vcells.net/dendritic-cell.

## ACKNOWLEDGEMENTS

We would like to thank Mr. Fitsumbirhan Mehari for his analysis on preliminary data of the project. The contribution of each author is described below using contributor roles taxonomy.

## Conceptualization

XL, JV; Data curation: XL, MC, ME, TJ, SU, CL; Formal analysis: XL; Funding acquisition: XL, JV; Investigation: XL, FD, KFG, SU, AE, JD, NS; Methodology: XL; Project administration: XL; Resources: XL, FD, AE, JW, HMJ, JD, NS, JV; Software: XL, MC, TJ; Supervision: XL, JV; Validation: XL, ME; Visualization: XL; Writing-original draft: XL, FD, MC, ME, JV; Writing-review & editing: XL, ME, JD, NS, JV.

## REFERENCE

1. Banchereau, J. and Steinman, R.M. (1998) Dendritic cells and the control of immunity. Nature, 392, 245–252.

2. Dörrie, J., Schaft, N., Schuler, G. and Schuler-Thurner, B. (2020) Therapeutic Cancer Vaccination with Ex Vivo RNA-Transfected Dendritic Cells-An Update. Pharmaceutics, 12.

3. Steinman, R.M. and Hemmi, H. (2006) Dendritic cells: translating innate to adaptive immunity. Curr. Top. Microbiol. Immunol., 311, 17–58.

4. Constantino, J., Gomes, C., Falcão, A., Neves, B.M. and Cruz, M.T. (2017) Dendritic cell-based immunotherapy: a basic review and recent advances. Immunol. Res., 65, 798–810.

5. Tacken, P.J., de Vries, I.J.M., Torensma, R. and Figdor, C.G. (2007) Dendritic-cell immunotherapy: from ex vivo loading to in vivo targeting. Nat. Rev. Immunol., 7, 790–802.

6. Hayden, M.S. and Ghosh, S. (2011) NF-κB in immunobiology. Cell Res., 21, 223–244.

7. Pfeiffer, I.A., Hoyer, S., Gerer, K.F., Voll, R.E., Knippertz, I., Gückel, E., Schuler, G., Schaft, N. and Dörrie, J. (2014) Triggering of NF-κB in cytokine-matured human DCs generates superior DCs for T-cell priming in cancer immunotherapy. Eur. J. Immunol., 44, 3413–3428.

8. Liu, T., Zhang, L., Joo, D. and Sun, S.-C. (2017) NF-κB signaling in inflammation. Signal Transduct Target Ther, 2.

9. Taniguchi, K. and Karin, M. (2018) NF-κB, inflammation, immunity and cancer: coming of age. Nat. Rev. Immunol., 18, 309–324.

10. Bartel, D.P. (2009) MicroRNAs: target recognition and regulatory functions. Cell, 136, 215–233.

11. Bartel, D.P. (2018) Metazoan MicroRNAs. Cell, 173, 20–51.

12. Turner, M.L., Schnorfeil, F.M. and Brocker, T. (2011) MicroRNAs regulate dendritic cell differentiation and function. J. Immunol., 187, 3911–3917.

13. Zhou, H. and Wu, L. (2017) The development and function of dendritic cell populations and their regulation by miRNAs. Protein Cell, 8, 501–513.

14. Inui, M., Martello, G. and Piccolo, S. (2010) MicroRNA control of signal transduction. Nat. Rev. Mol. Cell Biol., 11, 252–263.

15. Smyth, L.A., Boardman, D.A., Tung, S.L., Lechler, R. and Lombardi, G. (2015) MicroRNAs affect dendritic cell function and phenotype. Immunology, 144, 197–205.

16. Lu, C., Huang, X., Zhang, X., Roensch, K., Cao, Q., Nakayama, K.I., Blazar, B.R., Zeng, Y. and Zhou, X. (2011) miR-221 and miR-155 regulate human dendritic cell development, apoptosis, and IL-12 production through targeting of p27kip1, KPC1, and SOCS-1. Blood, 117, 4293–4303.

17. Lai, X., Schmitz, U., Gupta, S.K., Bhattacharya, A., Kunz, M., Wolkenhauer, O. and Vera, J. (2012) Computational analysis of target hub gene repression regulated by multiple and cooperative miRNAs. Nucleic Acids Res., 40, 8818–8834.

18. Lai, X., Gupta, S.K., Schmitz, U., Marquardt, S., Knoll, S., Spitschak, A., Wolkenhauer, O., Pützer, B.M. and Vera, J. (2018) MiR-205-5p and miR-342-3p cooperate in the repression of the E2F1 transcription factor in the context of anticancer chemotherapy resistance. Theranostics, 8, 1106–1120.

19. Lai, X., Eberhardt, M., Schmitz, U. and Vera, J. (2019) Systems biology-based investigation of cooperating microRNAs as monotherapy or adjuvant therapy in cancer. Nucleic Acids Res, 47, 7753–7766.

20. Hao Shi, null, Yan, K.-K., Ding, L., Qian, C., Chi, H. and Yu, J. (2020) Network Approaches for Dissecting the Immune System. iScience, 23, 101354.

21. Szeto, G.L. and Finley, S.D. (2019) Integrative Approaches to Cancer Immunotherapy. Trends Cancer, 5, 400–410.

22. Schaft, N., Dörrie, J., Thumann, P., Beck, V.E., Müller, I., Schultz, E.S., Kämpgen, E., Dieckmann, D. and Schuler, G. (2005) Generation of an optimized polyvalent monocyte-derived dendritic cell vaccine by transfecting defined RNAs after rather than before maturation. Journal of Immunology, 174, 3087–3097.

23. Gerer, K.F., Erdmann, M., Hadrup, S.R., Lyngaa, R., Martin, L.-M., Voll, R.E., Schuler-Thurner, B., Schuler, G., Schaft, N., Hoyer, S., et al. (2017) Preclinical evaluation of NF-κB-triggered dendritic cells expressing the viral oncogenic driver of Merkel cell carcinoma for therapeutic vaccination. Ther Adv Med Oncol, 9, 451–464.

24. Bosch, N.C., Voll, R.E., Voskens, C.J., Gross, S., Seliger, B., Schuler, G., Schaft, N. and Dörrie, J. (2019) NF-κB activation triggers NK-cell stimulation by monocyte-derived dendritic cells. Ther Adv Med Oncol, 11, 1758835919891622.

25. Hoyer, S., Gerer, K.F., Pfeiffer, I.A., Prommersberger, S., Höfflin, S., Jaitly, T., Beltrame, L., Cavalieri, D., Schuler, G., Vera, J., et al. (2015) Electroporated Antigen-Encoding mRNA Is Not a Danger Signal to Human Mature Monocyte-Derived Dendritic Cells. J Immunol Res, 2015, 952184.

26. Andrews, S. (2017) Babraham Bioinformatics - FastQC A Quality Control tool for High Throughput Sequence Data.

27. Li, H. (2013) Aligning sequence reads, clone sequences and assembly contigs with BWA-MEM. 1303.3997 [q-bio].

28. Kozomara, A. and Griffiths-Jones, S. (2014) miRBase: annotating high confidence microRNAs using deep sequencing data. Nucleic Acids Research, 42, D68–73.

29. Bourgon, R., Gentleman, R. and Huber, W. (2010) Independent filtering increases detection power for high-throughput experiments. Proc. Natl. Acad. Sci. U.S.A., 107, 9546–9551.

30. Tam, S., Tsao, M.-S. and McPherson, J.D. (2015) Optimization of miRNA-seq data preprocessing. Brief. Bioinformatics, 16, 950–963.

31. Love, M.I., Huber, W. and Anders, S. (2014) Moderated estimation of fold change and dispersion for RNA-seq data with DESeq2. Genome Biol., 15, 550.

32. R Core Team (2017) R: The R Project for Statistical Computing.

33. Benjamini, Y. and Hochberg, Y. (1995) Controlling the False Discovery Rate: A Practical and Powerful Approach to Multiple Testing. Journal of the Royal Statistical Society: Series B (Methodological), 57, 289–300.

34. Fabregat, A., Jupe, S., Matthews, L., Sidiropoulos, K., Gillespie, M., Garapati, P., Haw, R., Jassal, B., Korninger, F., May, B., et al. (2018) The Reactome Pathway Knowledgebase. Nucleic Acids Res., 46, D649–D655.

35. Wu, D. and Smyth, G.K. (2012) Camera: a competitive gene set test accounting for inter-gene correlation. Nucleic Acids Res, 40, e133–e133.

36. Ritchie, M.E., Phipson, B., Wu, D., Hu, Y., Law, C.W., Shi, W. and Smyth, G.K. (2015) limma powers differential expression analyses for RNA-sequencing and microarray studies. Nucleic Acids Res., 43, e47.

37. Wu, G., Feng, X. and Stein, L. (2010) A human functional protein interaction network and its application to cancer data analysis. Genome Biol., 11, R53.

38. Agarwal, V., Bell, G.W., Nam, J.-W. and Bartel, D.P. (2015) Predicting effective microRNA target sites in mammalian mRNAs. eLife, 4, e05005.

39. Chou, C.-H., Shrestha, S., Yang, C.-D., Chang, N.-W., Lin, Y.-L., Liao, K.-W., Huang, W.-C., Sun, T.-H., Tu, S.-J., Lee, W.-H., et al. (2018) miRTarBase update 2018: a resource for experimentally validated microRNA-target interactions. Nucleic Acids Res., 46, D296–D302.

40. Li, J.-H., Liu, S., Zhou, H., Qu, L.-H. and Yang, J.-H. (2014) starBase v2.0: decoding miRNA-ceRNA, miRNA-ncRNA and protein-RNA interaction networks from large-scale CLIP-Seq data. Nucleic Acids Res., 42, D92–97.

41. Bhattacharya, S., Dunn, P., Thomas, C.G., Smith, B., Schaefer, H., Chen, J., Hu, Z., Zalocusky, K.A., Shankar, R.D., Shen-Orr, S.S., et al. (2018) ImmPort, toward repurposing of open access immunological assay data for translational and clinical research. Sci Data, 5, 180015.

42. Schmeier, S., Alam, T., Essack, M. and Bajic, V.B. (2017) TcoF-DB v2: update of the database of human and mouse transcription co-factors and transcription factor interactions. Nucleic Acids Res., 45, D145–D150.

43. Cornish, A.J. and Markowetz, F. (2014) SANTA: quantifying the functional content of molecular networks. PLoS Comput. Biol., 10, e1003808.

44. Schmitz, U., Lai, X., Winter, F., Wolkenhauer, O., Vera, J. and Gupta, S.K. (2014) Cooperative gene regulation by microRNA pairs and their identification using a computational workflow. Nucleic Acids Res., 42, 7539–7552.

45. Pedersen, T.L. (2020) ggraph: An Implementation of Grammar of Graphics for Graphs and Networks.

46. Csardi, G. and Nepusz, T. (2015) igraph – Network analysis software.

47. Gu, Z., Eils, R. and Schlesner, M. (2016) Complex heatmaps reveal patterns and correlations in multidimensional genomic data. Bioinformatics, 32, 2847–2849.

48. Wickham, H. (2016) ggplot2: Elegant Graphics for Data Analysis 2nd ed. Springer International Publishing.

49. Allaire, J.J., Ellis, P., Gandrud, C., Kuo, K., Lewis, B.W., Owen, J., Russell, K., Rogers, J., Sese, C. and Yetman, C.J. (2017) networkD3: D3 JavaScript Network Graphs from R.

50. Shannon, P., Markiel, A., Ozier, O., Baliga, N.S., Wang, J.T., Ramage, D., Amin, N., Schwikowski, B. and Ideker, T. (2003) Cytoscape: a software environment for integrated models of biomolecular interaction networks. Genome Res., 13, 2498–2504.

51. Wentker, P., Eberhardt, M., Dreyer, F.S., Bertrams, W., Cantone, M., Griss, K., Schmeck, B. and Vera, J. (2017) An Interactive Macrophage Signal Transduction Map Facilitates Comparative Analyses of High-Throughput Data. J. Immunol., 198, 2191–2201.

52. Ma, X., Becker Buscaglia, L.E., Barker, J.R. and Li, Y. (2011) MicroRNAs in NF-kappaB signaling. J Mol Cell Biol, 3, 159–166.

53. Jin, P., Han, T.H., Ren, J., Saunders, S., Wang, E., Marincola, F.M. and Stroncek, D.F. (2010) Molecular signatures of maturing dendritic cells: implications for testing the quality of dendritic cell therapies. J Transl Med, 8, 4.

54. Gebert, L.F.R. and MacRae, I.J. (2019) Regulation of microRNA function in animals. Nat. Rev. Mol. Cell Biol., 20, 21–37.

55. Tan, H., Huang, S., Zhang, Z., Qian, X., Sun, P. and Zhou, X. (2019) Pan-cancer analysis on microRNA-associated gene activation. EBioMedicine, 43, 82–97.

56. Summers deLuca, L. and Gommerman, J.L. (2012) Fine-tuning of dendritic cell biology by the TNF superfamily. Nat. Rev. Immunol., 12, 339–351.

57. Locy, H., Melhaoui, S., Maenhout, S.K. and KrisThielemans (2018) Dendritic Cells: The Tools for Cancer Treatment. In.

58. Ji, R.-R., Chasalow, S.D., Wang, L., Hamid, O., Schmidt, H., Cogswell, J., Alaparthy, S., Berman, D., Jure-Kunkel, M., Siemers, N.O., et al. (2012) An immune-active tumor microenvironment favors clinical response to ipilimumab. Cancer Immunol. Immunother., 61, 1019–1031.

59. Comerford, I., Bunting, M., Fenix, K., Haylock-Jacobs, S., Litchfield, W., Harata-Lee, Y., Turvey, M., Brazzatti, J., Gregor, C., Nguyen, P., et al. (2010) An immune paradox: how can the same chemokine axis regulate both immune tolerance and activation?: CCR6/CCL20: a chemokine axis balancing immunological tolerance and inflammation in autoimmune disease. Bioessays, 32, 1067–1076.

60. Lee, A.Y.S., Eri, R., Lyons, A.B., Grimm, M.C. and Korner, H. (2013) CC Chemokine Ligand 20 and Its Cognate Receptor CCR6 in Mucosal T Cell Immunology and Inflammatory Bowel Disease: Odd Couple or Axis of Evil? Front Immunol, 4, 194.

61. Xiao, Z., Casey, K.A., Jameson, S.C., Curtsinger, J.M. and Mescher, M.F. (2009) Programming for CD8 T cell memory development requires IL-12 or type I IFN. J. Immunol., 182, 2786–2794.

62. Olsson Åkefeldt, S., Maisse, C., Belot, A., Mazzorana, M., Salvatore, G., Bissay, N., Jurdic, P., Aricò, M., Rabourdin-Combe, C., Henter, J.-I., et al. (2013) Chemoresistance of human monocyte-derived dendritic cells is regulated by IL-17A. PLoS ONE, 8, e56865.

63. Carrington, E.M., Zhang, J.-G., Sutherland, R.M., Vikstrom, I.B., Brady, J.L., Soo, P., Vremec, D., Allison, C., Lee, E.F., Fairlie, W.D., et al. (2015) Prosurvival Bcl-2 family members reveal a distinct apoptotic identity between conventional and plasmacytoid dendritic cells. Proc. Natl. Acad. Sci. U.S.A., 112, 4044–4049.

64. Gao, Y., Nish, S.A., Jiang, R., Hou, L., Licona-Limón, P., Weinstein, J.S., Zhao, H. and Medzhitov, R. (2013) Control of T helper 2 responses by transcription factor IRF4-dependent dendritic cells. Immunity, 39, 722–732.

65. Moynagh, P.N. (2005) The NF-kappaB pathway. J. Cell. Sci., 118, 4589–4592.

66. Hoesel, B. and Schmid, J.A. (2013) The complexity of NF-κB signaling in inflammation and cancer. Mol. Cancer, 12, 86.

67. Paardekooper, L.M., Bendix, M.B., Ottria, A., de Haer, L.W., Ter Beest, M., Radstake, T.R.D.J., Marut, W. and van den Bogaart, G. (2018) Hypoxia potentiates monocyte-derived dendritic cells for release of tumor necrosis factor α via MAP3K8. Biosci. Rep., 38.

68. Hegde, V.L., Singh, N.P., Nagarkatti, P.S. and Nagarkatti, M. (2008) CD44 mobilization in allogeneic dendritic cell-T cell immunological synapse plays a key role in T cell activation. J. Leukoc. Biol., 84, 134–142.

69. Termeer, C., Averbeck, M., Hara, H., Eibel, H., Herrlich, P., Sleeman, J. and Simon, J.C. (2003) Targeting dendritic cells with CD44 monoclonal antibodies selectively inhibits the proliferation of naive CD4+ T-helper cells by induction of FAS-independent T-cell apoptosis. Immunology, 109, 32–40.

70. Luan, Y.-Y., Yao, R.-Q., Tong, S., Dong, N., Sheng, Z.-Y. and Yao, Y.-M. (2016) Effect of tumor necrosis factor-α induced protein 8 like-2 on immune function of dendritic cells in mice following acute insults. Oncotarget, 7, 30178–30192.

71. Kabashima, K., Shiraishi, N., Sugita, K., Mori, T., Onoue, A., Kobayashi, M., Sakabe, J.-I., Yoshiki, R., Tamamura, H., Fujii, N., et al. (2007) CXCL12-CXCR4 engagement is required for migration of cutaneous dendritic cells. Am. J. Pathol., 171, 1249–1257.

72. Benvenuti, F., Hugues, S., Walmsley, M., Ruf, S., Fetler, L., Popoff, M., Tybulewicz, V.L.J. and Amigorena, S. (2004) Requirement of Rac1 and Rac2 expression by mature dendritic cells for T cell priming. Science, 305, 1150–1153.

73. Zanoni, I. and Granucci, F. (2012) Regulation and dysregulation of innate immunity by NFAT signaling downstream of pattern recognition receptors (PRRs). Eur. J. Immunol., 42, 1924–1931.

74. Bao, M., Wang, Y., Liu, Y., Shi, P., Lu, H., Sha, W., Weng, L., Hanabuchi, S., Qin, J., Plumas, J., et al. (2016) NFATC3 promotes IRF7 transcriptional activity in plasmacy--toid dendritic cells. J. Exp. Med., 213, 2383–2398.

75. Collins, L.E., DeCourcey, J., Rochfort, K.D., Kristek, M. and Loscher, C.E. (2014) A role for syntaxin 3 in the secretion of IL-6 from dendritic cells following activation of toll-like receptors. Front Immunol, 5, 691.

76. Gottipati, S., Rao, N.L. and Fung-Leung, W.-P. (2008) IRAK1: a critical signaling mediator of innate immunity. Cell. Signal., 20, 269–276.

77. McNab, F., Mayer-Barber, K., Sher, A., Wack, A. and O’Garra, A. (2015) Type I interferons in infectious disease. Nat. Rev. Immunol., 15, 87–103.

78. Finotti, G., Tamassia, N. and Cassatella, M.A. (2017) Interferon-λs and Plasmacytoid Dendritic Cells: A Close Relationship. Front Immunol, 8, 1015.

79. Anguille, S., Lion, E., Van den Bergh, J., Van Acker, H.H., Willemen, Y., Smits, E.L., Van Tendeloo, V.F. and Berneman, Z.N. (2013) Interleukin-15 dendritic cells as vaccine candidates for cancer immunotherapy. Hum Vaccin Immunother, 9, 1956–1961.

80. Mookerjee, A., Graciotti, M. and Kandalaft, L.E. (2019) IL-15 and a Two-Step Maturation Process Improve Bone Marrow-Derived Dendritic Cell Cancer Vaccine. Cancers (Basel), 11.

81. Wilcox, R.A., Chapoval, A.I., Gorski, K.S., Otsuji, M., Shin, T., Flies, D.B., Tamada, K., Mittler, R.S., Tsuchiya, H., Pardoll, D.M., et al. (2002) Cutting edge: Expression of functional CD137 receptor by dendritic cells. J. Immunol., 168, 4262–4267.

82. Lagali, N.S., Badian, R.A., Liu, X., Feldreich, T.R., Ärnlöv, J., Utheim, T.P., Dahlin, L.B. and Rolandsson, O. (2018) Dendritic cell maturation in the corneal epithelium with onset of type 2 diabetes is associated with tumor necrosis factor receptor superfamily member 9. Sci Rep, 8, 14248.

83. Shin, V.Y., Jin, H., Ng, E.K.O., Cheng, A.S.L., Chong, W.W.S., Wong, C.Y.P., Leung, W.K., Sung, J.J.Y. and Chu, K.-M. (2011) NF-κB targets miR-16 and miR-21 in gastric cancer: involvement of prostaglandin E receptors. Carcinogenesis, 32, 240–245.

84. Pla, A., Zhong, X. and Rayner, S. (2018) miRAW: A deep learning-based approach to predict microRNA targets by analyzing whole microRNA transcripts. PLoS Comput. Biol., 14, e1006185.

85. Pinzón, N., Li, B., Martinez, L., Sergeeva, A., Presumey, J., Apparailly, F. and Seitz, H. (2017) microRNA target prediction programs predict many false positives. Genome Res., 27, 234–245.

86. Amon, L., Lehmann, C.H.K., Baranska, A., Schoen, J., Heger, L. and Dudziak, D. (2019) Transcriptional control of dendritic cell development and functions. Int Rev Cell Mol Biol, 349, 55–151.

87. Ye, H., Liu, X., Lv, M., Wu, Y., Kuang, S., Gong, J., Yuan, P., Zhong, Z., Li, Q., Jia, H., et al. (2012) MicroRNA and transcription factor co-regulatory network analysis reveals miR-19 inhibits CYLD in T-cell acute lymphoblastic leukemia. Nucleic Acids Res., 40, 5201–5214.

88. Bracken, C.P., Scott, H.S. and Goodall, G.J. (2016) A network-biology perspective of microRNA function and dysfunction in cancer. Nat. Rev. Genet., 17, 719–732.

89. Winkler, I., Bitter, C., Winkler, S., Weichenhan, D., Thavamani, A., Hengstler, J.G., Borkham-Kamphorst, E., Kohlbacher, O., Plass, C., Geffers, R., et al. (2020) Identification of Pparγ-modulated miRNA hubs that target the fibrotic tumor microenvironment. Proc. Natl. Acad. Sci. U.S.A., 117, 454–463.

90. Hayden, M.S., West, A.P. and Ghosh, S. (2006) NF-kappaB and the immune response. Oncogene, 25, 6758–6780.

91. Kanehisa, M., Furumichi, M., Tanabe, M., Sato, Y. and Morishima, K. (2017) KEGG: new perspectives on genomes, pathways, diseases and drugs. Nucleic Acids Res., 45, D353–D361.

92. Slenter, D.N., Kutmon, M., Hanspers, K., Riutta, A., Windsor, J., Nunes, N., Mélius, J., Cirillo, E., Coort, S.L., Digles, D., et al. (2018) WikiPathways: a multifaceted pathway database bridging metabolomics to other omics research. Nucleic Acids Res., 46, D661–D667.

93. Mubeen, S., Hoyt, C.T., Gemünd, A., Hofmann-Apitius, M., Fröhlich, H. and Domingo-Fernández, D. (2019) The Impact of Pathway Database Choice on Statistical Enrichment Analysis and Predictive Modeling. Front Genet, 10, 1203.

94. Geistlinger, L., Csaba, G., Santarelli, M., Ramos, M., Schiffer, L., Turaga, N., Law, C., Davis, S., Carey, V., Morgan, M., et al. (2020) Toward a gold standard for benchmarking gene set enrichment analysis. Brief. Bioinformatics, 10.1093/bib/bbz158.

95. Khan, F.M., Marquardt, S., Gupta, S.K., Knoll, S., Schmitz, U., Spitschak, A., Engelmann, D., Vera, J., Wolkenhauer, O. and Pützer, B.M. (2017) Unraveling a tumor type-specific regulatory core underlying E2F1-mediated epithelial-mesenchymal transition to predict receptor protein signatures. Nat Commun, 8, 198.

96. Dreyer, F.S., Cantone, M., Eberhardt, M., Jaitly, T., Walter, L., Wittmann, J., Gupta, S.K., Khan, F.M., Wolkenhauer, O., Pützer, B.M., et al. (2018) A web platform for the network analysis of high-throughput data in melanoma and its use to investigate mechanisms of resistance to anti-PD1 immunotherapy. Biochim. Biophys. Acta, 1864, 2315–2328.

97. Moreau, Y. and Tranchevent, L.-C. (2012) Computational tools for prioritizing candidate genes: boosting disease gene discovery. Nat. Rev. Genet., 13, 523–536.

98. Guala, D. and Sonnhammer, E.L.L. (2017) A large-scale benchmark of gene prioritization methods. Sci Rep, 7, 46598.

99. Iván, G. and Grolmusz, V. (2011) When the Web meets the cell: using personalized PageRank for analyzing protein interaction networks. Bioinformatics, 27, 405–407.

100. Hsu, C.-L., Huang, Y.-H., Hsu, C.-T. and Yang, U.-C. (2011) Prioritizing disease candidate genes by a gene interconnectedness-based approach. BMC Genomics, 12 Suppl 3, S25.

101. McGeary, S.E., Lin, K.S., Shi, C.Y., Pham, T.M., Bisaria, N., Kelley, G.M. and Bartel, D.P. (2019) The biochemical basis of microRNA targeting efficacy. Science, 366.

